# Dual-site high-density 4Hz transcranial alternating current stimulation applied over auditory and motor cortical speech areas does not influence auditory-motor mapping

**DOI:** 10.1101/514687

**Authors:** Basil Preisig, Matthias J. Sjerps, Anne Kösem, Lars Riecke

## Abstract

**Background:** Verbal repetition of auditory speech requires the mapping of a sensory acoustic input onto articulatory motor plans (auditory-motor mapping). Recent evidence indicates that auditory-motor mapping could rely on low frequency neural synchronization (i.e., theta oscillatory phase coupling) between sensory and motor speech areas.

**Objective:** In the present study, we apply dual-site high-density (HD) Transcranial Alternating Current Stimulation (TACS) above the auditory and motor cortex to induce, or disrupt, theta phase coupling between the two areas. We predicted that functionally coupling the two areas would strengthen auditory-motor mapping, compared with functionally decoupling them. We assessed the strength of auditory-motor mapping using a verbal repetition task.

**Results:** We found no significant effect of TACS-induced theta phase coupling on auditory-motor mapping as indexed by verbal repetition performance.

**Conclusion:** Auditory-motor mapping may rely on a different mechanism than we hypothesized, for example, oscillatory phase-coupling outside the theta range. Alternatively, modulation of interregional theta-phase coupling may require more effective stimulation protocols, for example, TACS at higher intensities.

## 1 Introduction

Speaking requires the mapping of speech sounds onto articulatory motor plans (auditory-motor mapping). According to current speech processing models, the mapping of auditory speech representations onto articulatory motor plans is accomplished by the dorsal processing stream. This processing stream connects the posterior superior temporal lobe and the ventral premotor cortex/posterior Broca’s area via a sensorimotor interface in the Sylvian fissure at the boundary between the parietal and temporal lobes (area SPT) [1–3]. There is substantial evidence for dorsal stream involvement in auditory-motor mapping [4–8]. However, the neural bases of auditory-motor mapping are only partially understood.

Previous research in healthy participants and in patients with brain lesion has provided causal evidence for dorsal stream processing during auditory-motor mapping: For instance, it has been shown that auditory-motor mapping, as measured by verbal repetition of auditory sentences, pseudowords, or syllables, can be modulated in healthy participants by exciting or inhibiting cortical regions belonging to the left dorsal stream with Transcranial Magnetic Stimulation (TMS) [5,6]. Moreover, studies using various methods (histology, lesion mapping and diffusion tensor-imaging) have provided converging evidence for the relevance of dorsal stream integrity for auditory-motor mapping [4,7–9].

In the present study, we further investigated the mechanism underlying the interaction (or ‘communication’) between auditory and motor cortex during auditory-motor mapping. We hypothesized that the communication between the two regions is mediated by interregional synchronization of local theta (3-7Hz) oscillations [10–12]. This idea is motivated by two recent studies that have investigated frontotemporal oscillatory coupling during passive story listening [13] and syllable processing [14]. Park and colleagues [13] found that the frontal and motor cortices modulate the phase of speech-coupled low-frequency oscillations in the auditory cortex. Moreover, Assaneo & Poeppel [14] reported that neural oscillatory synchronization of auditory and motor cortex during syllable processing is restricted to the theta range, peaking at ∼4.5Hz, close to the natural syllable rate across languages [14]. More generally, the theta frequency band is thought to play a functional role in auditory speech perception because it overlaps with intelligibility-relevant temporal fluctuations in the acoustic speech signal (∼1-8Hz). Ongoing theta oscillations in auditory cortex can align to the amplitude envelope of an acoustic speech signal, a phenomenon called speech-brain entrainment [15] that may contribute to the linguistic/phonological analysis of speech signals (for a review see [16]).

Previous brain stimulation studies have shown that speech perception can be modulated with 4-Hz transcranial alternating current stimulation (TACS), suggesting a causal role of theta speech-brain entrainment in speech comprehension [17]. Here, we used TACS to disrupt or enhance the communication between auditory and motor areas during auditory-motor mapping, by modulating interregional phase-coupling of local theta oscillations in auditory cortex and motor cortex. We assessed the causal influence of TACS-induced interregional phase-coupling on auditory-motor mapping using a behavioral task that required participants to listen to and verbally repeat nonwords. Verbal repetition is a auditory-motor task commonly used to assess dorsal stream processing, i.e., auditory-motor mapping [2,18]. To induce interregional theta phase-coupling (synchronization) between auditory cortex and motor cortex, we applied theta (4Hz) TACS simultaneously above each of the two regions (in-phase stimulation). Theta-TACS is thought to entrain theta oscillatory phase in cortex ([19–22], but see also [23]). To decouple (desynchronize) the two regions, we included a second TACS condition in which we reversed the phase of TACS above one of the regions (anti-phase stimulation). We predicted that stronger auditory-motor theta phase-coupling (in-phase stimulation vs. anti-phase stimulation) would lead to an improvement in auditory-motor mapping that would be reflected by an improvement in verbal repetition performance.

To allow for a manipulation check, we further varied the phase relationship between the auditory syllables and the 4-Hz TACS. We reasoned that the observation of a cyclic modulation of verbal repetition performance would validate our a priori assumption that TACS entrained theta oscillatory phase, which is difficult to verify with simultaneously obtained electrophysiological measures [24]. Our results reveal no significant effect of our theta-phase coupling manipulation on auditory-motor mapping.

## 2 Material & Methods

### 2.1 Participants

19 right-handed healthy volunteers (13 female, age: *M* = 22, *SD* = 2.33) participated in the study. All participants had normal or corrected-to-normal visual acuity. The participants reported no history of neurological, psychiatric, or hearing disorders. They had normal hearing (defined as hearing thresholds of less than 25 dB HL at 250, 500, 750, 1000, 1500, 3000, and 4000Hz), according to a pure tone audiometry screening. All participants fulfilled the inclusion criteria for noninvasive brain stimulation as assessed by a prior screening and gave written informed consent prior to the experiment. Ethical approval to conduct this study was provided by the local ethics committee (CMO region Arnhem-Nijmegen). The present study was conducted in accordance with the principles of the latest version of the Declaration of Helsinki.

### 2.2 Stimuli

Trisyllabic nonword stimuli were created out of a sequence of three Dutch consonant vowel (CV) syllables. CVs were composed of the voiceless consonants, /k/, /p/, and /t/, and the vowels /y/, /u/, and /y/ (“uu”, “oe” and “uh”). This led to a total of 9 combinations that were recorded from a female Dutch speaker. The vowel part of the recorded CVs was time-compressed using Praat [25] to obtain natural sounding syllables, but with equal durations (125ms). Syllables were interleaved with 125-ms silent gaps, resulting in a syllable rate of 4Hz within each nonword stimulus. The nonwords were built based on all CV combinations that contained three different consonants and three different vowels, which led to a set of 36 unique stimuli. A random subset of 18 stimuli was used for training and main experiment in the first session, and the other half in the second session. All nonword stimuli were presented in stationary noise that was matched to the average spectrum of the speech.

### 2.3 Electric stimulation

Electric currents were applied through two HD electrode configurations each consisting of concentric conductive rubber electrodes [26]: a central round electrode, radius = 1.0 cm, and a surrounding ring electrode (inner radius = 3.5 cm, outer radius = 4.0 cm). The central electrodes were placed according to the international 10-20 system, in between FT7 and FC5 (frontal configuration), and P7 and P5 (temporal configuration). These configurations were chosen to produce relatively strong currents in the target regions, which were the speech motor areas (i.e., left inferior frontal cortex) and auditory speech areas (i.e., left superior temporal cortex), as suggested by prior electric field simulations on a standard head model using the simnibs toolbox [27].

Sinusoidal current stimulation was applied through two battery-driven transcranial current stimulators (Neuroconn, Ilmenau, Germany). The frequency of the TACS current matched the syllable rate of the nonword stimuli, i.e., 4Hz. TACS intensity was set to 1mA peak-to-peak and kept constant across participants. The current density was 0.4 mA/cm^2^ at the center and 0.1 mA/cm^2^ at the concentric ring electrode. Impedance was kept below 10 kΩ. Before starting the actual experiment, we assured that all participants well tolerated stimulation intensity. Stimulation was ramped over the first and the last 10 s of each experimental block using raised-cosine ramps.

The timing of the electric and auditory stimuli was controlled using a multichannel D/A converter (National Instruments, sampling rate: 16kHz) and Datastreamer software [28]. Visual stimulation and response recording were controlled using Presentation^®^ software (Version 18.0, Neurobehavioral Systems, Inc., Berkeley, CA).

### 2.4 Experimental design and task

Verbal repetition performance was assessed using a verbal repetition task that required participants to listen to each nonword and to verbally repeat it as accurately and as quickly as possible. The verbal response intervals were defined by inter-stimulus interval (ISI) of approximately 6 seconds. The exact ISI depended on the phase lag condition of two consecutive trials. The experimental procedure included two sessions that were conducted within 5 to 10 days. Participants were seated in a sound-attenuated booth and first familiarized with the verbal repetition task in a stepwise procedure: First, they listened to and repeated all CV syllables three times. The noise in this phase of the experiment was set to a moderate level (SNR: +10 dB). Second, nonword repetition was practiced audiovisually (i.e., with the addition that written nonwords were presented on the screen) at the same noise level. Third, nonword repetition was practiced again without visual presentation. Finally, to avoid potential range effects, an adaptive staircase procedure [29] was used that identified the individual noise level (SNR, *M* = −0.64 *SD* = 3.04) yielding an intermediate performance level of 70%.

Each session of the experiment consisted of three experimental blocks each representing a different stimulation condition: *in-phase, anti-phase*, or *sham stimulation*. 1) *In-phase stimulation* was applied with relative phase lag of 0° between the central electrodes placed over the motor (i.e., left inferior frontal) and sensory speech areas (i.e., left superior temporal lobe), i.e., frontotemporal synchronization. 2) *Anti-phase stimulation* was applied with a relative phase lag of 180° between the central frontal electrode relative to the central temporal electrode, i.e., frontotemporal desynchronization. 3.) During *sham stimulation* (placebo) the onset ramp was followed immediately by an offset ramp, i.e., no stimulation was applied during the actual experiment. The ramp was repeated at the end of the block. The order of stimulation conditions was reversed across consecutive sessions and counterbalanced across participants.

Following the procedure of previous studies [17,20,21], the relative timing of TACS and nonword stimuli was manipulated across six different phase-lag conditions by varying the onset of the nonword stimuli in steps of 30° (41.7ms) across the 4Hz TACS cycle. For a visual illustration of the different experimental conditions see Figure 1.

**Figure 1.**
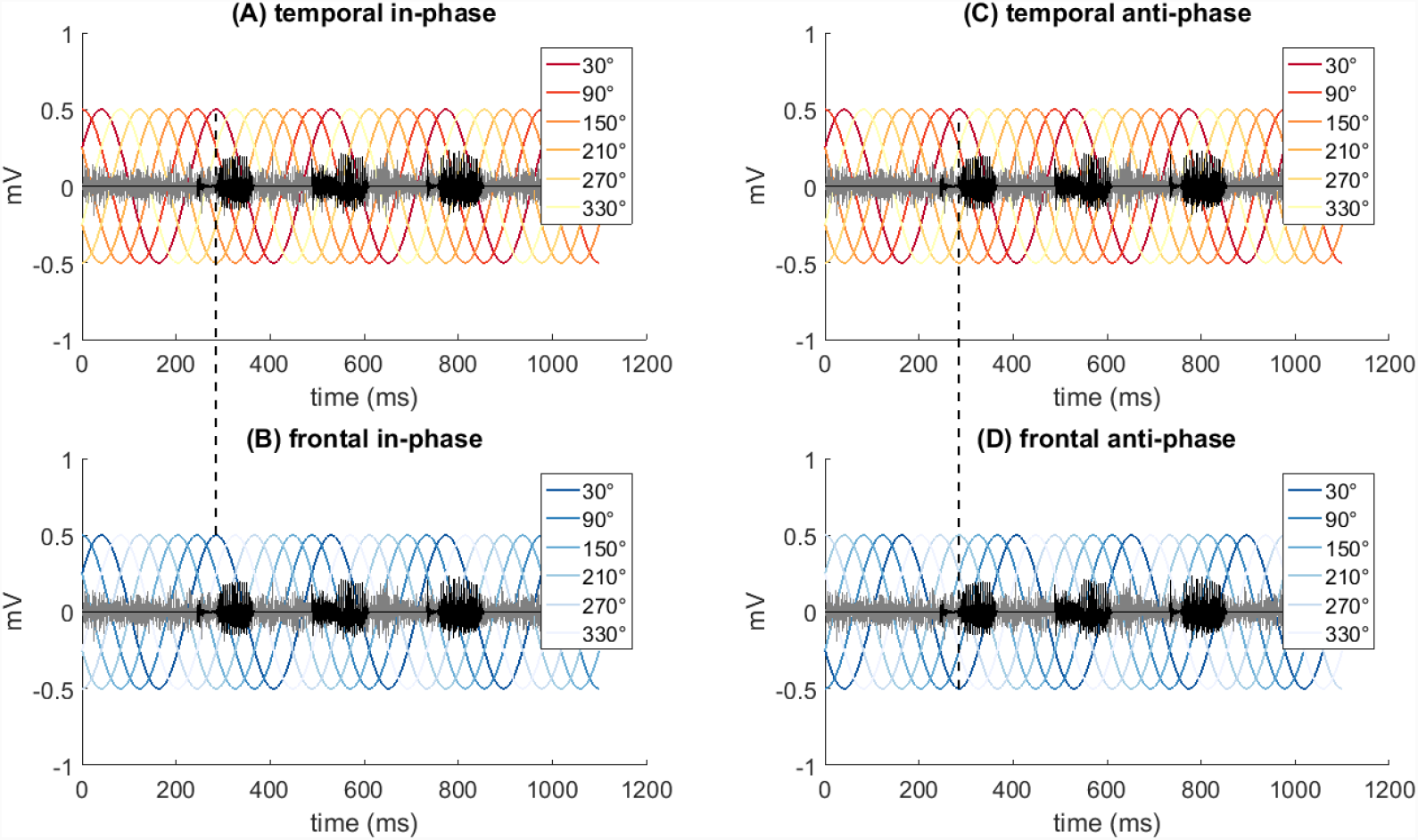
Synchronization of electric and auditory stimulation, and synchronization/desynchronization of temporal and frontal cortex. This is an illustration of the timing between the sinusoidal electric currents (chromatic colors), the sound pressure curve of the stimulus (black) and the frequency matched noise (grey). Sinusoids with different chromatic colors represent the six phase lag conditions. In-phase stimulation (A+B) was applied with a relative phase lag of 0° (dotted line) between the temporal (A) and the frontal cortex (B), i.e., frontotemporal synchronization. Anti-phase stimulation (C+D) was applied with a relative phase lag of 180° (dotted line) between the temporal (C) relative to the frontal cortex (D), i.e., frontotemporal desynchronization.

### 2.5 Data analysis

The behavioral data, i.e., recorded participant responses, were analyzed as follows: First, a blinded research assistant scored the recordings from all trials offline. Afterwards, author BCP, who was also blinded, scored one randomly selected block per participant. Interrater agreement was high (raw agreement: 97% and Cohen’s kappa: 0.85). Second, repetition performance was calculated in each condition as the percentage of correctly repeated trials. Third, for each participant and each stimulation condition, the six phase-lag conditions were concatenated to build a time series that enabled assessing periodic variations in repetition performance across the TACS cycle.

The phase lag that leads to the best performance (best lag) may vary across participants because of anatomical differences, i.e., the relative orientation of the current flow to the stimulated neural tissue. To compensate for such potential inter-individual differences, the phase-lag series was realigned to the best lag (defined here as 0°) under the assumption that TACS modulated performance at the critical 4Hz frequency. Importantly, to avoid non-independency the best lag was excluded from subsequent analyses, as it necessarily represents the maximum of the time series due to the best-phase alignment.

We tested whether interregional theta-phase coupling modulated behavioral performance by comparing the aligned time series in the in-phase condition with the aligned time series in the anti-phase condition. We further verified whether theta oscillatory phase indeed entrained to our 4Hz TACS by comparing phase lags near the best lag (distance from best lag: 60°; −60°), which are supposed to delimit an ‘excitatory’ 4Hz half-cycle associated with relatively good behavioral performance, with more distant lags (distance from best lag: 120°; −240°) supposed to delimit the opposite, i.e., an ‘inhibitory’ half-cycle associated with poorer performance. As an additional test, we calculated the spectral density of the behavioral time series by applying the Fast Fourier transformation and compared the power of the resulting different frequency components within and across the different stimulation conditions (in-phase; anti-phase; sham). If 4Hz TACS effects were caused by entrainment of theta oscillations, we expected behavior to modulate in the theta range, and hence to observe maximum power at the 4Hz component.

Statistical testing was done using parametric tests for repeated measures. ANOVAs were used to test for stimulation effect, session effect, phase (or frequency) effect, and interactions. Post hoc comparisons were done using paired t-tests.

## 3 Results

Participants correctly responded on average 73.67% ± 3.5% (mean ± SEM) of the trials. The average best lag across participants and stimulation conditions was 99° ± 23° (mean ± SEM), which is equivalent to an audio lag of 69.05ms ± 10.61ms (mean ± SEM). The distribution of the participants’ best lags pooled across TACS conditions (in-phase and anti-phase) did not deviate significantly from uniformity (*z* = 1.789, *p* = 0.168), suggesting that best lag varied across participants. Moreover, best lag did not correlate significantly between sessions or stimulation conditions (*ps* >.70). Therefore, participants’ data were aligned to the best lag separately for each session and stimulation condition.

To test our main hypothesis regarding interregional phase coupling, we analyzed the impact of Stimulation condition across phase lags and sessions. A three-way repeated measures ANOVA, including the within subject-factors *Stimulation condition* (in-phase; anti-phase; sham), *Phase lag* condition after alignment (−60°; 60°; 120°; 180°; 240°), and *Session* (1; 2) was conducted. The analysis did not reveal a main effect of *Session* (*p* >.11), nor an interaction *Stimulation condition* x *Phase lag* x *Session* (*p*>.39). Therefore, the data were pooled across sessions and a two-way repeated measures ANOVA, including only *Stimulation condition* (in-phase; anti-phase; sham) and *Phase lag* (−60°; 60°; 120°; 180°; 240°) as factors was conducted. Contrary to our predictions, we found no interaction *Stimulation condition* x *Phase lag* (*p* >.57), and no significant main effect of *Stimulation condition* (*p* >.91) (Figure 2).

**Figure 2.**
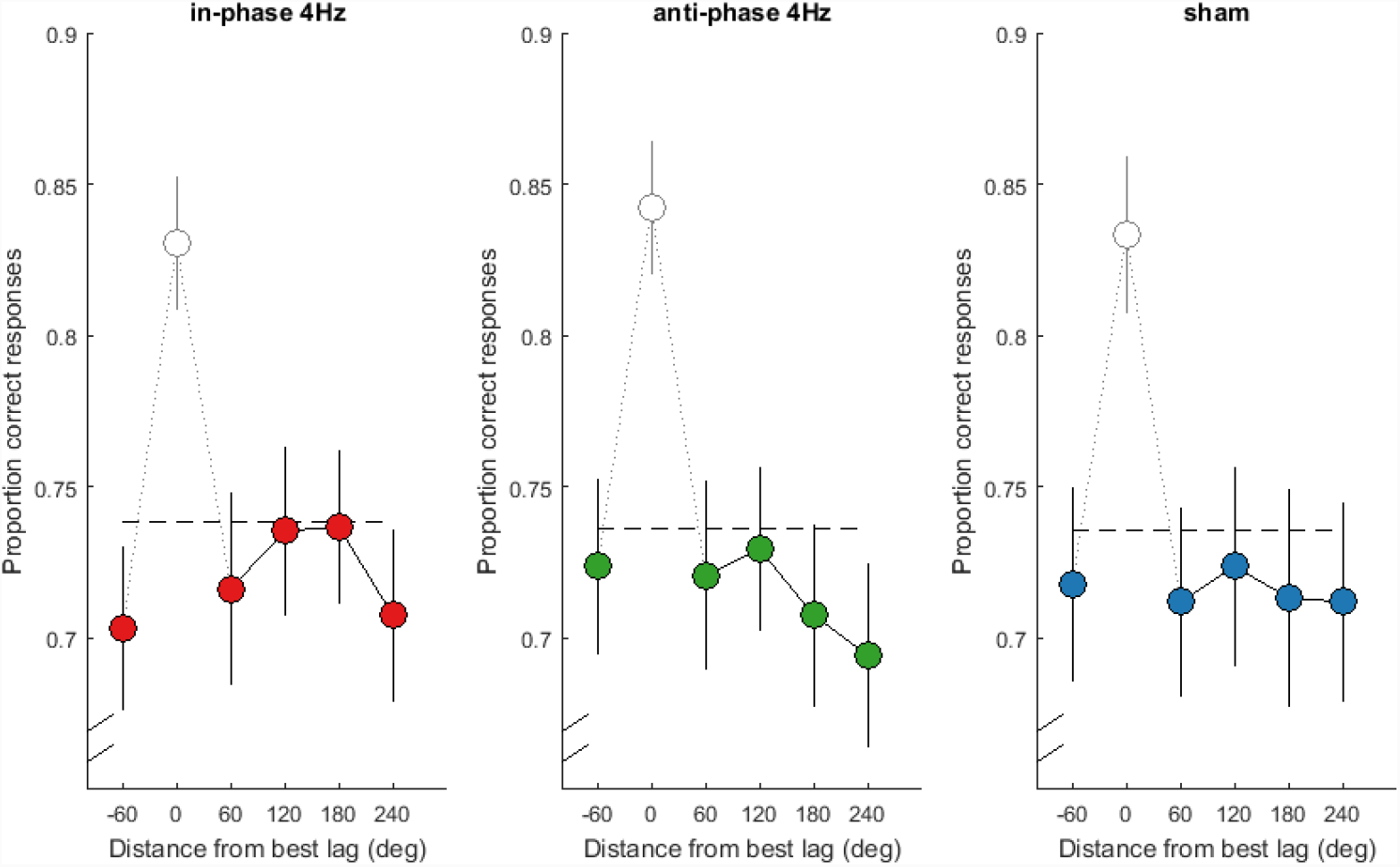
Participants’ average performance (mean ± SEM across participants) as a function of phase lag condition is shown for each stimulation condition (in-phase 4Hz; anti-phase 4Hz, and sham). The peak performance at the best lag (0°) is trivial and was excluded from the analysis. The horizontal line represents average performance per stimulation condition. Contrary to our predictions, performance (pooled across phase-lag conditions) did not differ significantly across the three stimulation conditions.

To verify whether TACS entrained local theta oscillations independent of auditory-motor mapping, we assessed whether performance was better at the presumed excitatory 4Hz half cycle (−60°; 60°) vs. the presumed inhibitory half cycle (120°; 240°). A two-way repeated measures ANOVA including the within-subject factors *TACS half cycle* (excitatory; inhibitory) and *Stimulation Condition* (TACS; sham, the former was pooled across the in- and anti-phase condition), revealed no significant interaction or main effect of *TACS half cycle* (*p* >.69) (Figure 3A). As an additional test of TACS phase, the spectral density of the behavioral time series was analyzed. A two-way repeated measures ANOVA including the factors *Frequency* (4Hz; 8Hz; 12Hz) and *Stimulation condition* (TACS; sham) was conducted. The analysis revealed a significant main effect of *Frequency* [*F*(2,36) = 43.12, *p* < .01, *η2* = 0.388], but no interaction *Stimulation condition* x *Frequency* (*p* >.32), nor a main effect of *Stimulation condition* (*p* >.70). Spectral power was significantly lower at 12Hz compared to 4Hz and 8Hz (*ps* <.01, Holm-Bonferroni-corrected) (Figure 3B).

**Figure 3.**
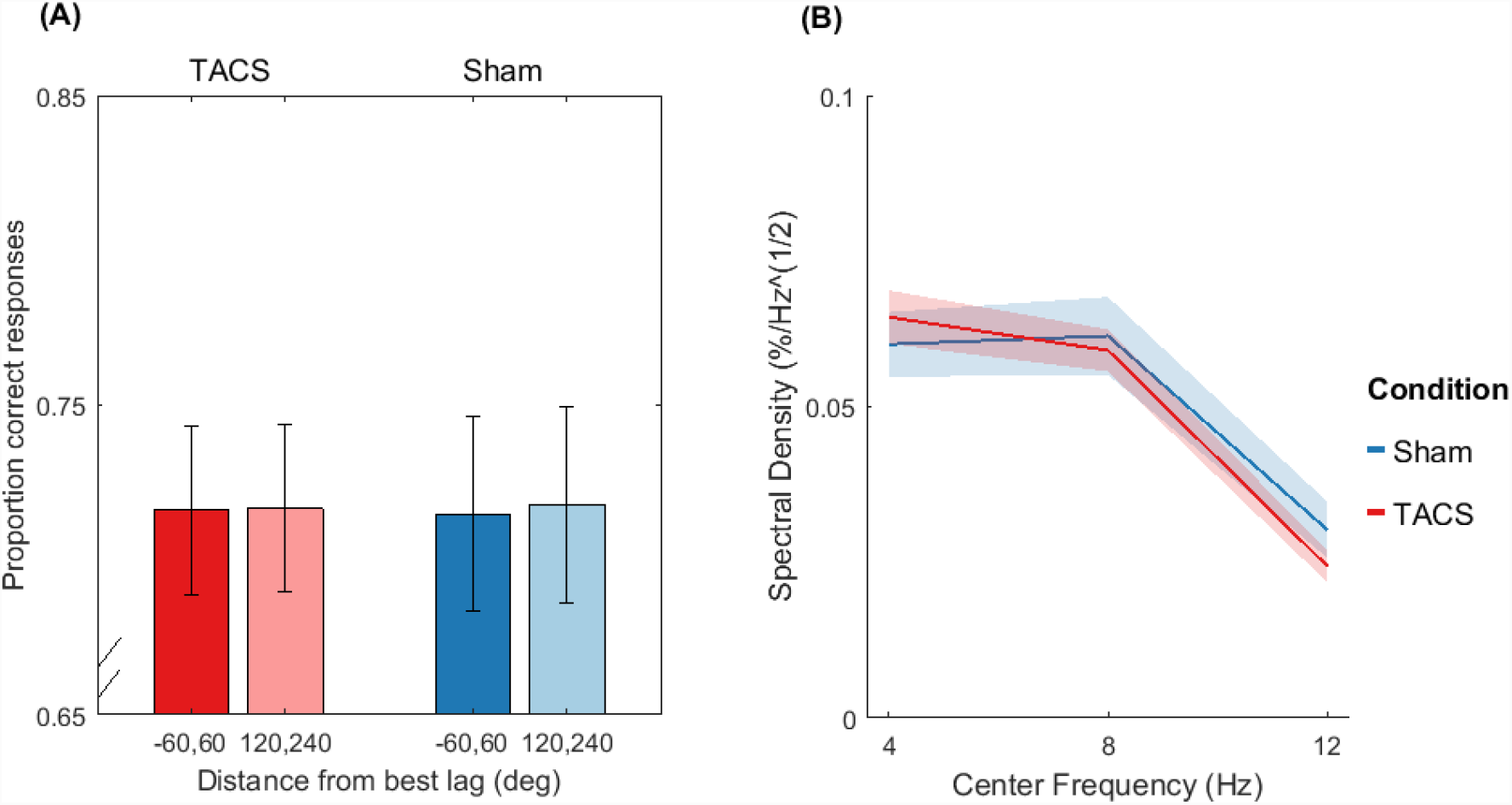
(A) The average performance (mean ± SEM across participants) across the best-lag distances presumed to resemble an excitatory half-cycle (−60, 60) and an inhibitory half-cycle (120, 240) did not differ between TACS (pooled in-phase and anti-phase) and sham. (B) The power spectrum of the behavioral times series (six phase lag conditions; see Figure 2) is shown for TACS vs sham. Our prediction was that power peaks at the component corresponding to the TACS frequency and decreases monotonically at higher frequencies. The analysis revealed indeed a main effect of frequency and significantly lower power at 12Hz compared to 4Hz and 8Hz (*ps* <.01, Holm-Bonferroni-corrected), but these effects were not significantly stronger under TACS vs. sham.

In addition, we conducted exploratory analyses involving the application of the above analysis to alternative measures (duration and latency of verbal productions). These analyses, which were not in the focus of the study design, led to similar null results.

## 4 Discussion

The aim of this study was to test whether inducing or disrupting the functional coupling between auditory and motor speech areas in the theta range modulates auditory-motor mapping. We hypothesized that theta phase coupling between the two areas would strengthen auditory-motor speech mapping and that this would be observable as an improvement in listeners’ ability to verbally repeat nonwords. To test this, we applied theta-TACS simultaneously above listeners’ auditory cortex and motor cortex (either in-phase or anti-phase) to synchronize or desynchronize the two areas and we measured the effect on verbal repetition performance. To verify whether TACS successfully entrained theta oscillations, we varied the relative phase of the auditory stimuli and TACS.

Our results show no significant effect of stimulation condition, i.e., no significant difference in performance between in-phase compared to anti-phase stimulation, and no significant difference between real stimulation compared to sham stimulation. In contrast to previous studies on the auditory perception threshold [20,21,30] and speech comprehension [17,31], we found no significant periodic variation in performance across TACS phases.

Our results indicate that auditory-motor mapping may rely on mechanisms different from auditory-motor theta phase-coupling. Alternatively, it may rely on auditory-motor theta phase-coupling as we have hypothesized, but we failed to observe this because of potential methodological shortcomings. In the following paragraphs, we will discuss the two potential interpretations in more detail.

First, auditory-motor mapping may rely on mechanisms different from auditory-motor theta phase coupling. For example, auditory-motor mapping might depend on oscillatory phase coupling, but in frequency bands outside the theta range that we investigated. The latter idea is supported by two studies; for example, Schoffelen et al. [32] found that the direction of information flow between language-relevant brain areas depends on the contribution of distinct frequency bands while participants were visually presented with word lists and sentences. They found that rhythmic activity in the alpha frequency range (8–12Hz) propagates from temporal cortical areas to frontal cortical areas, whereas beta activity (15–30Hz) propagates in the opposite direction, when participants read sentences and word lists during MEG recording. Moreover, the results by Park and colleagues [13] indicate that top-down communication from the left inferior frontal gyrus to the left auditory cortex during speech perception may be stronger in the delta frequency band than the theta frequency band. Future studies may investigate effects of phase-coupling in the delta, alpha, or beta range on auditory-motor mapping using delta, alpha, or beta TACS respectively.

A second potential interpretation is that our verbal repetition task was not sensitive enough to capture the presumed variations in auditory-motor mapping strength. However, Murakami and colleagues [6] found TMS-induced modulations of auditory-motor mapping applying a similar verbal repetition task. Their participants had to repeat single syllables and pseudowords embedded in white noise. Furthermore, Restle et al [33] found TMS-induced effects while participants had to repeat sentences in a foreign language. These previous findings thus indicate that our behavioral measure (verbal repetition performance) is suited to capture effects of non-invasive brain stimulation.

Third, in contrast to previously applied TMS protocols [6,33], perhaps our theta-TACS protocol was not effective enough to modulate verbal auditory-motor mapping strength. This could be related to two main parameters: electrode placement and stimulation intensity. For conventional electrode configurations, larger electrodes (standard size 5 x 7 cm) are usually placed over bilateral homologue stimulation sites, which leads to an extended electric field spanning the area between the two stimulation electrodes in both cerebral hemispheres. In contrast, unilateral HD configurations, like the concentric configuration applied in the current study, usually induce more focal electric fields that are more restricted to the region of interest and surrounding brain tissue in the hemisphere under the electrodes. The improved focality comes at the cost of a lower current quantity penetrating the brain; because of the smaller distance between the electrodes, more current is shunted through the skull or the cerebrospinal fluid [34]. Maybe this disadvantage of unilateral HD-TACS outweighed the theoretical advantage of higher focality [35] in the current study. Concerning stimulation intensity, it must be acknowledged that the stimulation intensity in the present study (1mA peak-to-peak) was lower than the average stimulation intensity in studies showing effects of theta-TACS on auditory perception (1.6mA ±0.1 peak-to-peak) and speech comprehension (1.8mA ±0.1 peak-to-peak) [17,20,21]. The reason for the lower stimulation intensity was that participants’ sensation threshold tends to be lower with the HD-configuration due to the relatively high current density related to the smaller electrodes. In line with this potential limitation, it has been shown that transcranial direct current stimulation is more effective in enhancing cortical excitability when applied at 2mA than at 1mA ([36], but see also [37]). Finally, there currently is controversy about whether theta-TACS at its conventional intensity (1-2mA peak-to-peak) is powerful enough to entrain neural oscillations. Lafon and colleagues [23] found no reliable effect of extracortical theta-TACS on intracranial theta activity measured from implanted grid-electrodes in epilepsy patients.

Fourth, our TACS protocol may have been in fact effective (i.e., it entrained theta oscillatory phase), but the motor act and/or the sound accompanying the utterance caused a reset of theta phase that destroyed the TACS-induced phase entrainment. Indeed, a key difference to studies that found theta-TACS-induced modulations of working memory [11,38–41] may be that our participants had to perceive and verbally reproduce the presented nonwords. Moreover, compared with TACS speech studies that used verbal repetition to test the recognition of meaningful sentences [17,31], our speech stimuli were much shorter and thus provided less opportunity for oscillations to recover from any articulation-induced phase reset.

Related to the previous point, it might be that in the previous TACS speech studies, successful speech recognition required the participants to identify primarily the temporal structure of the speech stimuli, thus facilitating the chunking and prediction of linguistic elements. In contrast, successful verbal repetition of the artificial words in our study probably required more articulatory cues. TACS may have provided temporal, not articulatory, cues that helped our participants to identify the fixed temporal structure of our syllable sequences, not the identity of the individual syllables. However, if articulation indeed disrupts endogenous and/or exogenously induced theta entrainment, this would further question the functional role of theta-phase coupling in auditory-motor mapping. This is an interesting question that should be addressed in future studies. In this view, the role of theta oscillations is thus limited to the temporal analysis of speech and auditory speech perception, but not speech production/articulation.

In sum, based on these considerations the lack of an effect of theta phase coupling on auditory-motor mapping may be ascribed to different physiologic mechanisms, i.e., phase coupling in a different frequency band. Moreover, methodological limitation cannot be ruled out, specifically insufficient TACS intensity. These interpretations could be further tested in future studies by inducing interregional phase coupling within and across frequencies in the delta, alpha, or beta range with dual-site HD TACS at higher intensity

### 4.1 Conclusion

In sum, our results do not conclusively advocate for a functional role of theta phase coupling in auditory-motor mapping. Given the results of previous reports, neural phase coupling in other bands, e.g., delta, alpha, or beta may play such a role. The results further highlight potential limitations of unilateral dual-site HD TACS in interregional (de-) synchronization.

## 5 Acknowledgments

This work was supported by a grant awarded to B.C.P by the Swiss National Science Foundation [P2BEP3_168728]. The authors would like to thank Birgit Knudsen, Iris Schmidt, and Michel-Pierre Jansen for their assistance.

